# HERV-K and HERV-W transcriptional activity in Myalgic Encephalomyelitis/ Chronic Fatigue Syndrome

**DOI:** 10.1101/693465

**Authors:** Lucas S Rodrigues, Luiz H da Silva Nali, Cibele O D Leal, Ester C Sabino, Eliana M Lacerda, Caroline C Kingdon, Luis Nacul, Camila M Romano

## Abstract

Chronic Fatigue Syndrome / Myalgic Encephalomyelitis (CFS / MS) is an incapacitating chronic disease that dramatically compromise the life quality. The CFS/ME pathogenesis is multifactorial, and it is believed that immunological, metabolic and environmental factors play a role. It is well documented an increased activity of Human endogenous retroviruses (HERVs) from different families in autoimmune and neurological diseases, making these elements good candidates for biomarkers or even triggers for such diseases. Here the expression of Endogenous retroviruses K and W (HERV–K and HERV–W) was determined in blood from moderately and severely affected ME/CFS patients. HERV-K was overexpressed only in moderately affected individuals and HERV-W showed no difference. This is the first report about HERV-K differential expression in moderate ME/CFS.

## Introduction

Myalgic Encephalomyelitis/ Chronic Fatigue Syndrome (ME/CFS) is a chronic and debilitating disease with unknown etiology [1]. Affected individuals have compromised motor and cognitive capacities. There is a wide variation in the symptoms of this disease, which include joint pains, mood disturbance, and malaise and worsening of symptoms following minimal physical or mental exertion. More severe symptoms can be also present including extreme exhaustion, severe joint pains with no apparent cause, non-restorative sleep and a range of immune and neurological symptoms. These symptoms may lead to depression and social isolation in the person with ME/CFS [1]. The pathophysiology of the ME/CFS is not understood and there is no diagnostic biomarker available. There is still controversy over the etiology of the disease; however, it is widely accepted that several immunological alterations are present in ME/CFS patients [2]. In addition, accumulated evidence for an association of ME/CFS with viral infections also exists and many patients report the onset of their symptoms during or right after a flu-like illness [3]. Thereafter, an unusual autoimmune response against the infection would be responsible for the perpetuation of the ME/CFS symptoms. Viral participation is finally supported by the evidences of clinical benefit of patients treated with valganciclovir [4]. Unfortunately, the absence of large cohort studies that investigate at the molecular level the participation of infectious agents on the ME/CFS pathogenesis impairs our understanding of this disease.

Human Endogenous Retroviruses (HERVs) are derived from exogenous retroviral infections, which occurred early in the evolution of vertebrates. Due to active replication and transposition events, HERVs are extensively distributed through the host genome and constitute about 8% of the human genome [5]. Due to accumulated mutations over the primate and human evolution, most HERVs are non-functional, but intact open reading frames of some HERVs persist and can be reactivated in response to systemic and environmental factors such as hormones, stress, and infection by exogenous viruses including almost all human herpesviruses, HIV and others [6,7]. Given their potential pathogenic effects, which include molecular mimicry and immune deregulation, HERVs are often postulated as possible causes of autoimmune diseases. Among the more than 30 families, the K and W families are the most recently integrated, the most active, and have been frequently associated with neurological and autoimmune diseases such as multiple sclerosis, diabetes mellitus SLE, ALS and rheumatoid arthritis [8].

To our knowledge, only two studies have investigated the participation of endogenous retroviruses in ME/CFS with contrasting results [9,10].

Given the extensively described altered patterns of HERVs in several diseases and the gap in knowledge of its expression in ME/CFS, we investigated the expression of the HERVs K and W in patients diagnosed with ME/CFS.

## Methods

### Participants

We used PBMC samples from a hundred patients diagnosed with ME/CFS and stored in the UK ME/CFS Biobank (UKMEB) at the London School of Hygiene and Tropical Medicine in this study. The UKMEB is among the few biorepositories worldwide with advanced storage and linked research infrastructure dedicated to research into ME/CFS [11]. Seventy five samples were requested from participants diagnosed with moderate fatigue (ME/CFSm), and 25 from participants with severe fatigue (ME/CFSs). Participants with ME/CFS were defined as moderate or severely affected based on their mobility: those described as severely affected were house-bound or bed-bound, while those described as having mild/moderate ME/CFS were ambulatory [11]. Samples from 70 healthy controls (also provided by the UKMEB) were included. This study was approved by LSHTM and University of São Paulo ethical committees [#EC.2017.02 and #2728254 respectively].

### RNA extraction and Real Time PCR

RNA extraction from the PBMC samples was performed by the Trizol-chloroform method, with 1ml Trizol and subsequent addition of chloroform to solubilize lipids allowing its removal. The samples were centrifuged at 15,000 rpm for 15 minutes and the upper phase containing the RNA was further used. The material was precipitated with Isopropanol 100% and washed with 75% Ethanol. In both steps the material was centrifuged at 15,000 RPM for 10 minutes at 4 °C. After this process, the pellets were dried at room temperature for 10 minutes, and the RNA was eluted in 45μl of Nuclease-Free H2O. The decontamination of remnant DNA was performed using two rounds DNAse treatment (Turbo DNA-Free (Ambion) following the manufacturer’s instruction. The absence of DNA was confirmed by real time PCR without reverse transcriptase using primers for HERV-K or HERV-W (see primers description below). After this procedure, cDNA was synthesized using the High capacity cDNA Reverse Transcription kit (Ambion, USA) according the manufacturer’s instructions. Real-time PCRs were performed for the HERV-W, - K and the endogenous gene using the primers and conditions used previously by Nali et al [12] using the Sybr Green method. The primers used are described in Table 1.

**Table 1.**
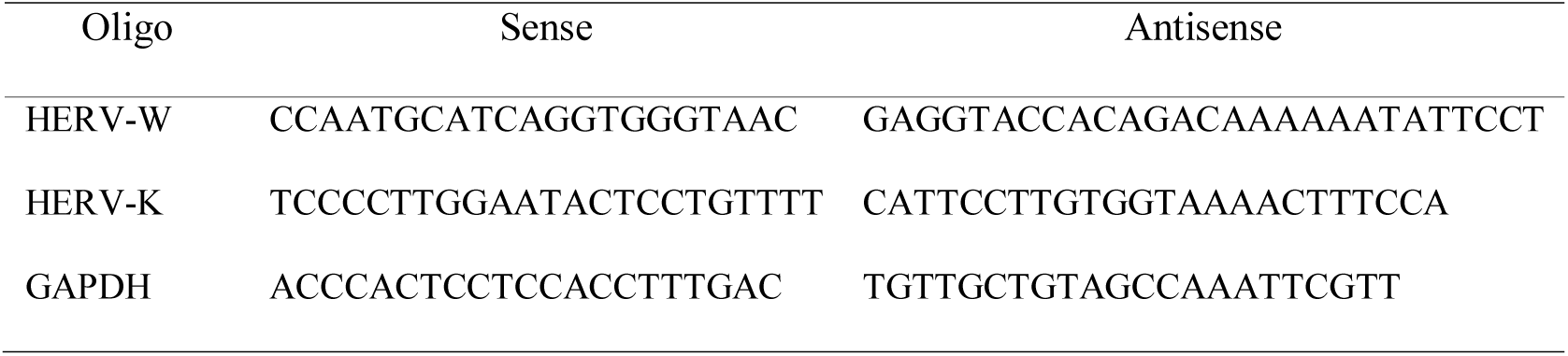
Primers used in real-time PCR assays

The cycling conditions for both HERVs detection were: 50 °C for 2 minutes, 95 °C for 10 minutes, followed by 40 cycles of 95 °C for 15 seconds, 50 °C for 1 minute, 60 °C for 1 minute. HERV activity was qualitatively (referred as presence/absence) and quantitatively (level of expression) evaluated. As positive controls we used a plasmid containing both HERV-W envelope and HERV-K polymerase fragments correspondent to the region covered by the primers used. The level of expression was determined by calculation of 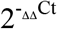, and the results were represented as fold changes. Statistical analysis was performed using the Wilcox test in the GraphPad Prism program v.6.04. Samples were only considered positive for HERVs and included in the analyses if expression of the endogenous control was also detected.

## Results

General description of individuals included in the study is described in Table 2. As expected, women were 4 times more prevalent than men. Therefore, we adjusted the control group to the same gender prevalence. HERV-K and W expression were evaluated in ME/CFS patients and healthy controls; and some level of expression of HERV-W was detected in all patients with severe fatigue and in 72/75 ME/CFSm (96%). HERV-K was also detected in all severe cases but in 65/75 of moderate cases (86.6%). The healthy control group was very similar to the moderate group, with 68/70 (97%) and 60/70 (85.7%) presenting expression of HERV-W and HERV-K respectively (Table 2). Only one patient with moderate fatigue and one control individual had no HERV activity at all. No relation was observed regarding HERV detection and duration of disease.

**Table 2.** Main characteristics of individuals included in the study.

| Participants (#) | Age<br>*median (max, min)* | Gender (%) |  | Time of disease<br>*median (max, min)* | Detection of HERV activity* |  |
| --- | --- | --- | --- | --- | --- | --- |
|  |  | M | F |  | HK | HW |
| ME/CFSs (25) | 43.3 (25-62) | 24% | 76% | 16.8 (2.8-40) | 100% | 100% |
| ME/CFSm (75) | 22.9 (18-64) | 25,4% | 74,6% | 11.2 (0.2 – 33.7) | 86.6% | 96% |
| Controls (70) | 42.8 (19-63) | 25.7% | 74.3% | - | 85.7% | 97% |

\* Qualitatively

Regarding to the level of expression (quantitative analysis), real time results revealed that HERV-W did not present significant differences when the healthy controls (HCs) or the two ME/CFS groups were compared between each other (Figure 1 A), i.e. ME/CFSs vs HCs (p = 0.89), ME/CFSm vs HCs (p = 0.77), ME/CFSs vs ME/CFSm (p = 0.95), all patients ME/CFS vs HCs (p = 0.78).

**Figure 1.**
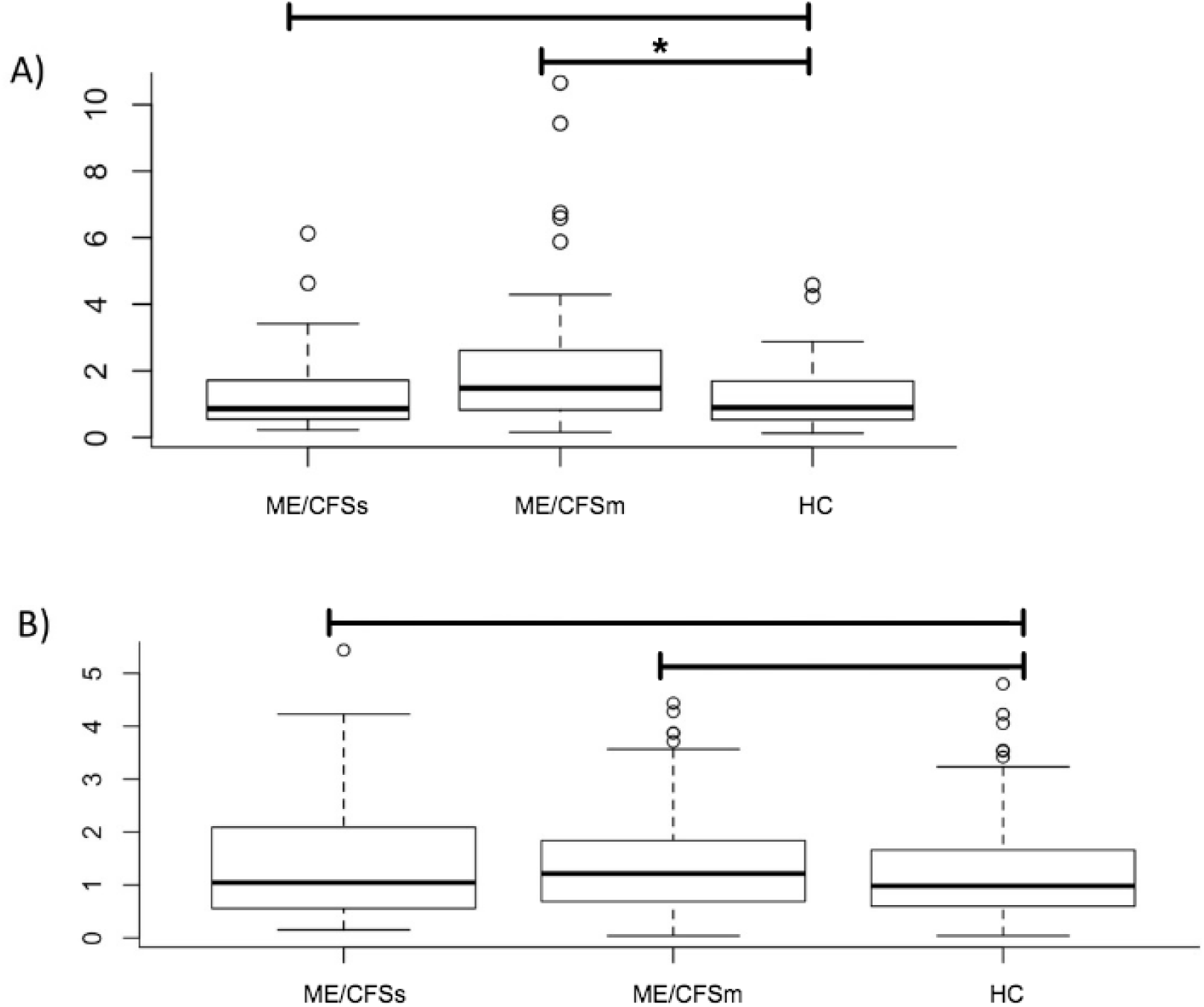
Boxplot of expression levels (in fold change) of HERVs among the groups. A) HERV-K and B) HERV-W. Significance between the groups (obtained by Wilcox test) is evidenced by an asterisk.

On the other hand, the HERV-K expression differed significantly between ME/CFSm group and the HCs (p = 0.050). HERV-K activity was not distinct between the ME/CFS groups: ME/CFSs vs ME/CFSm (p = 0.12), ME/CFS vs HCs (p = 0.17). ME/CFSs vs HCs (p = 0.97) (Figure 1 B).

## Discussion

The most recognized and widely-used case definitions (Fukuda [13] and Canadian Consensus criteria [1]) are based on self-reported symptoms. Studies of energy metabolism, oxidative stress and immunological alterations in ME/CFS have demonstrated imbalance in all these pathways, but the use of such information for diagnostic purposes is still far from reality.

Here, HERV-K and W transcripts were detected in all groups investigated, and we found that HERV-K was overexpressed in moderate ME/CFS. It is possible that the immunological, genic expression and metabolic alterations are different according to disease severity.

The interplay between endogenous retroviruses and the immune system is complex. ERVs are part of the host genome and in theory, they are supposed to be recognized as self-antigens and an immune tolerance should be established during the early stages of the organism development [14]. However, HERV products can interact with components of the innate immune system leading to the activation of pro-inflammatory pathways or, in some particular cases, their suppression [15]. The syncytin -2 protein for example, is a product of the ERV-FRD Env gene that has an immunosuppressive role by preventing maternal immune response against the fetus [16]. In a distinct scenario, it was demonstrated using psoriasis model that a pro-inflammatory environment could be able to suppress the expression of repetitive elements, including HERVs [17]. It would be reasonable to suggest that the immunological enhancement seen in more severe ME/CFS works by silencing the HERV transactivation that occurs in moderate cases. Such transactivation could be caused by exogenous viral replication or another as yet unknown factor. In line with this, Montoya and colleagues (2017) found a cytokine signature of severity in people with ME/CFS [18]. They demonstrated that from the 17 cytokines related to severity, 13 are pro-inflammatory, and (in addition to the worsening of the symptoms) may cause the reversion of the HERV-K activity to levels similar to those seen in healthy individuals. It may similarly occur with HERV-W, which, despite not being at significant levels, there was a slight decrease in people severely affected by ME/CFS when compared to those who are moderately affected.

Infection has often been considered as a trigger to ME/CFS. Many patients report that the fatigue began during or short after an episode of infectious disease. A number of pathogens including viruses have been associated with this disease [3]. And, due to its life long persistence and broad cell tropism, the herpesviridae family, particularly HHV-6 has been considered to be a possible trigger for ME/CFS for many, even though such relationship has not been consistent [3,9]. Interestingly, HHV-6 as some other herpesviruses, is also capable of transactivating HERVs, particularly, HERV-K [6]. Such transactivation may be either direct (through LTR activation by viral products) or indirect (via transcriptional binding factors and cytokines produced by viral replication) [3,6]. It is possible that as the disease progresses, whatever the exogenous infection that would have act as the trigger factor is controlled, and consequently, the HERVs transactivation decrease. Unfortunately, we did not perform serological or molecular tests for exogenous viruses.

Recently, it was suggested that differential methylation patterns of promoters in ME/CFS would impact on the expression of nearby transposable elements, including HERVs [19]. Two reports of HERV activity in ME/CFS were published some years ago but the results were conflicting. In 2013 Oakes and his team found no difference on the expression of HERV-K18 envelope in people with ME/CFS when compared with HCs [9]. In the same year De Meirleir and colleagues, using immunohistochemical methods, found immunoreactivity to HERV proteins (HERV-K, HERV-18, HERV-R and HERV-FRD) in dendritic cells of the duodenum of individuals diagnosed with the syndrome [10], suggesting that alterations in endogenous retroviruses expression pattern may occur in ME/CFS. The differences between the results of Oakes and colleagues and ours may be due to the methods used to detect HERV-K. While the present work used generic primers for HERV-K that allow the detection of hundreds of elements from most HML subfamilies the Oakes team searched for the HERV-K 18 envelope transcripts only, using a method specific to this particular element, while neglecting all the remaining proviruses from the K family. On the other hand, we are unable to determine which K family proviruses are involved in the differential expression observed.

The molecular method used here to detect HERV-W was also generic and was widely used in several studies that found differential expression of this element in pathological conditions, including in the blood, brain and CSF of multiple sclerosis (MS) patients [20]. Therefore, despite the similarity of a number of symptoms and the strong immunological component of ME/CFS and MS, the mechanisms responsible for HERV reactivation in such diseases are likely distinct.

In conclusion, this is the first report that demonstrates increased expression of an endogenous retrovirus in the blood of individuals with moderate ME/CFS. While the increased expression of these retroelements can’t be directly associated to the ME/CFS pathogeny, the observation of this phenomenon cannot be ignored.

